# Stable membrane topologies of small dual-topology membrane proteins

**DOI:** 10.1101/133298

**Authors:** Nir Fluman, Victor Tobiasson, Gunnar von Heijne

## Abstract

The topologies of α-helical membrane proteins are generally thought to be determined during their cotranslational insertion into the membrane. It is typically assumed that membrane topologies remain static after this process has ended. Recent findings, however, question this static view by suggesting that some parts of, or even the whole protein, can reorient in the membrane on a biologically relevant time scale. Here, we focus on anti-parallel homo-or hetero-dimeric Small Multidrug Resistance proteins, and examine whether the individual monomers can undergo reversible topological inversion (flip-flop) in the membrane until they are trapped in a fixed orientation by dimerization. By perturbing dimerization using various means, we show that the membrane topology of a monomer is unaffected by the presence or absence of its dimerization partner. Thus, membrane-inserted monomers attain their final topologies independently of dimerization, suggesting that wholesale topological inversion is an unlikely event *in vivo*.

## Introduction

α-Helical Membrane proteins make up about 25% of the proteomes of all living organisms (1) and mediate a plethora of essential functions. Despite their importance, our understanding of how they are produced to form folded, assembled, functional proteins *in vivo* is still quite limited.

The vast majority of membrane proteins are inserted into the membrane cotranslationally by ribosomes docked onto Sec-type translocons (2). Cotranslational insertion is a crucial decision-making step for membrane proteins, because achieving the correct membrane topology is essential for the protein’s ability to properly fold and function. Currently, the prevailing view is that topologies are determined cotranslationally and then remain fixed (3). This is because any topological change would require reorientation of hydrophobic transmembrane helices (TMHs) in the membrane and the transfer of hydrophilic loops through the membrane. However, although intuitively such processes are expected to be slow, several studies have provided evidence that topologies might not always remain static after the cotranslational insertion step. For example, the translocon initially misses two of the six TMHs of human aquaporin 1, which insert only later (4, 5). The transporter LacY from *E. coli* was shown to flip the topology of 7 out of its 12 TMHs upon manipulating the lipid composition of the membrane or upon phosphorylation, well after the protein has finished translation (6–8). Finally, the N-terminal TMH in the holin family of phage proteins inserts into the bacterial inner membrane only when the proton-motive force is reduced below a critical threshold (9).

Recent studies on EmrE from *E. coli,* a 4TMH multidrug transporter that belongs to the Small Multidrug Resistance (SMR) family (10), have suggested that it may also undergo topological rearrangements after the initial membrane-insertion event (11–14). The functional form of EmrE is a homodimer, in which the two identical protomers in the dimer have opposite membrane topologies and associate in antiparallel orientations relative to one another (15–19), although less stable parallel dimers can also form under certain conditions (16, 20–23). EmrE is the most extensively studied example of the class of dual-topology proteins: single polypeptide chains that are inserted into the membrane in two opposite orientations and form antiparallel homodimers *in vivo* (15, 24, 25). Curiously, the antiparallel dimers in the SMR family are not always homodimeric. In some family members the dimer is a heterodimer, composed of two non-identical but homologous subunits, each having a unique orientation opposite to its partner (24, 25).

The question of how a protein can be inserted with a precisely balanced 50:50 dual-topology is still incompletely understood. One of the factors involved is the positive-inside rule, which posits that membrane proteins orient such that loops containing more lysines and arginines face the cytosolic side of the membrane (26). In the single-topology, heterodimeric SMR proteins, the two monomers that form the dimer show strong and opposing biases in the distribution of lysines and arginines across the membrane (K/R-bias) (24). Dual-topology proteins such as EmrE lack a strong K/R-bias, resulting in a lack of strong preference towards one or the other orientation. What is not yet clear however, is if the weak K/R-bias is sufficient to generate the observed 50:50 topological distribution of EmrE protomers. One additional possibility that was recently proposed is post-translational dynamic reorientation of the monomer in the membrane (12, 13). This would be an elegant way to ensure a 50:50 distribution, since all monomers would eventually be trapped in antiparallel dimers, regardless of their initial topology.

Dynamic topological behavior of EmrE monomers was originally suggested based on the observation that a charged residue added to the very C-terminus of the protein can impact the orientation of the whole monomer (11). Further, it has recently been established that the N-terminal TMH of EmrE is initially inserted with both orientations in equal probabilities, likely before the C-terminus attains its final topology (13). Interestingly, biasing the topology of the C-terminus towards the cytosol (C_in_, for C-terminus inside the cell) by the addition of one or more positive charges causes the N-terminal TMH to reorient in the membrane (13). The topological change likely propagates through an aberrant “frustrated” 3-TMH topology to finally generate the fully inserted 4-TMH N_in_/C_in_ topology. Similar posttranslational topological dynamics were recently demonstrated also for the C-terminal TMH of EmrE. Again, when starting from an artificially generated, frustrated 3-TMH topology, the C-terminal TMH can flip in the membrane to generate a fully inserted 4-TMH topology (12). Indeed, simulations suggest that the topology even of wild type EmrE monomers initially inserted with frustrated topologies may be dynamic, with the different loops being transferred through the membrane with rates that are estimated to vary in the seconds-to-minutes range (14).

Notably, a recent study also suggested that fully inserted EmrE monomers with a 4-TMH topology can undergo dynamic flip-flop in the membrane (12). Here, we report a set of experiments to systematically test if SMR monomers are able to flip-flop freely in the membrane. We reasoned that if this were the case, then, as noted above, dimerization would serve as an energetic sink, trapping the monomers in the orientation needed for productive dimer association. We perturbed the dimerization of several model SMR proteins using different means and directly measured the effect on their membrane orientation. Our studies show that dimerization does not affect the balance between N_in_-C_in_ and N_out_-C_out_ topologies, implying that SMR monomers have stable topologies already before they dimerize. Any topological flip-flop in the SMR family of proteins is therefore probably confined to a brief instance immediately after synthesis, where after monomers remain in stable 4-TMH topologies that do not undergo wholesale flip-flop in the membrane, irrespective of subsequent dimerization events.

## Results

### Heterodimeric SMRs orient correctly in the membrane before dimerizing

As a first step towards studying topological dynamics in SMRs, we considered the possibility that their topologies might remain dynamic even after the cotranslational insertion step. If so, it might be expected that some SMR monomers be first inserted with an incorrect orientation, and then undergo flip-flop until they are trapped in a stable orientation by dimerization with a cognate partner. Since the membrane orientation is determined mainly via the positive inside rule (2, 26), we reasoned that any such initially mis-oriented subunits might have an insufficient lysine and arginine bias (K/R-bias) across the membrane. We therefore analyzed the K/R-biases in genomic pairs of SMR transporters – which are predicted to form anti-parallel heterodimers (24), where each protein subunit has a unique topology. We identified 224 SMR heterodimer genes that occur in pairs in putative gene operons (112 non-redundant pairs). We then predicted the locations of their four TMHs, and calculated their K/R-bias. Figure 1*A* shows that, as expected, in the vast majority of SMR pairs, one member has a strong negative K/R-bias and the other member has a strong positive one. Such strong signals would from the outset orient the two subunits in an anti-parallel fashion, as needed for forming the anti-parallel heterodimer. We did, however, find a few cases in which only one of the two members has a strong K/R-bias and the other partner only has a weak such bias (Fig. 1*A*, red circles, Fig. 1*B*). The weakly biased proteins could potentially be mis-oriented upon insertion and depend on dimerization with the more strongly biased partner to become locked in their correct orientation.

**Figure 1.**
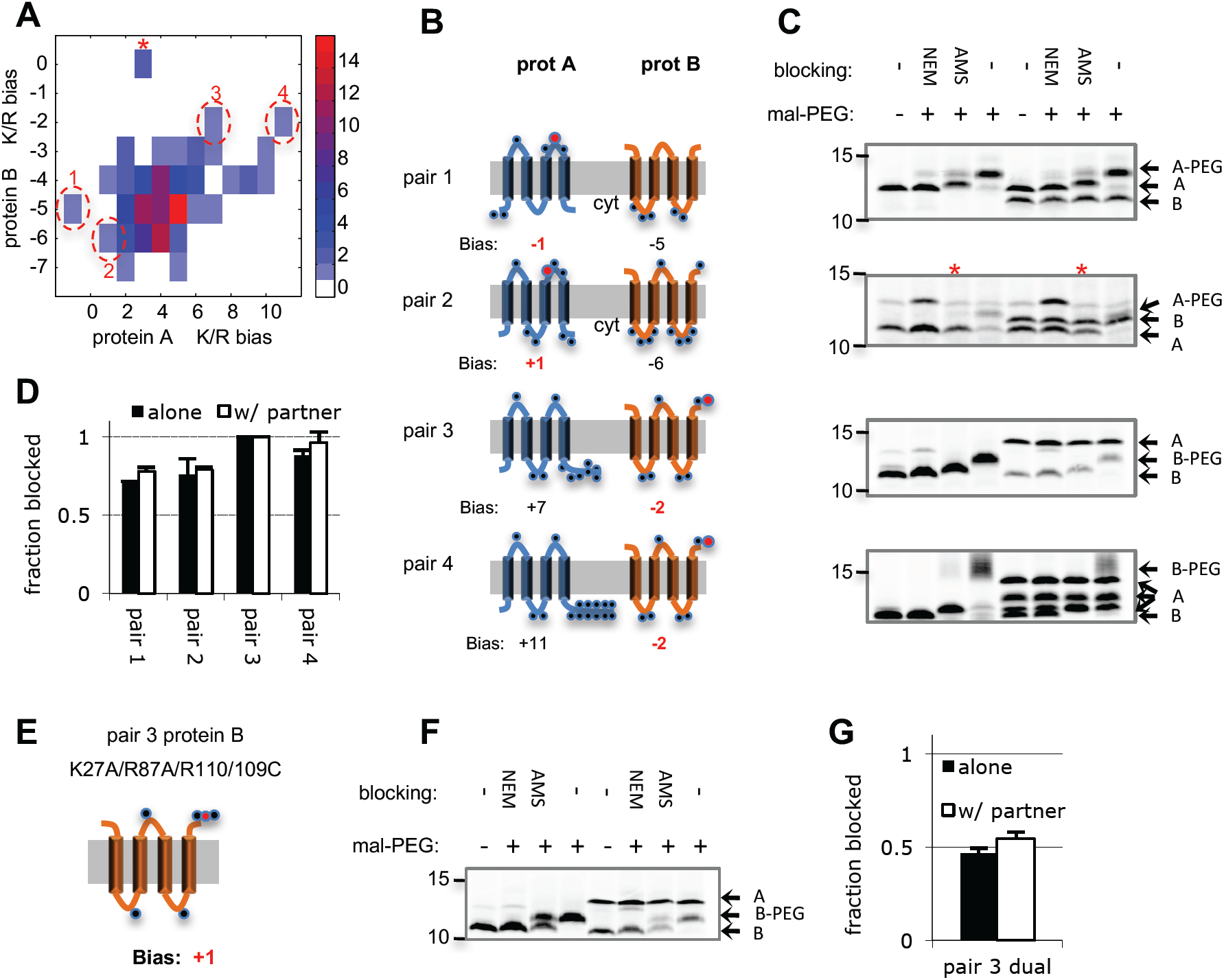
Analysis of heterodimeric SMR protein pairs. (A) Positive charge biases in heterodimeric SMR transporter pairs. Non-redundant heterodimeric SMR gene pairs were identified from genomic sequences (112 pairs, see Materials and Methods). The transmembrane positive charge (K/R) biases of the different pairs are shown in a heat map, where color indicates the number of observed occurrences of pairs with the indicated K/R-biases. The subunit with more positive bias was assigned as protein A, regardless of the genomic gene order. Genes pairs that were studied further are encircled in red and numbered. (*) Note that an interesting pair marked in asterisk was not studies further due to multiple cysteines in its sequence. (B) Schemes showing predicted topologies of the selected SMR pairs. The membrane is shown in gray, with the cytosolic side facing down. Black circles indicate positive charges (K or R) and the K/R-bias is indicated below the proteins, with the partner with weak bias in red. Red circle indicates the position of the introduced single cysteine mutation. (C) SCAM of single Cys mutants expressed with or without their genomic partners. The addition of various blockers to whole cells and of mal-PEG to solubilized membranes is given above. The first and last four lanes show proteins expressed alone or with their partners, respectively. Migration of molecular weight standards is given on the left. The migration of the different protein subunits and their PEGylated versions is shown by arrows. (*) Note that for pair #2, MTSET was used as a periplasmic blocker instead of AMS, see Fig. S1 for results using AMS and further explanation. (D) Quantification of SCAM results for the weakly biased members in all pairs, expressed alone or together with their partners. Error bars, SD of 3 independent experiments. (E) Model of dual topology version of pair 3 protein B. Compare the positive charges (black circles) with the same protein in section (*B*) of this figure. (F and G) SCAM results and their quantification, respectively, for the dual topology pair 3 protein B expressed alone or together with its partner.

To test this possibility, we generated single-Cys versions of the weakly biased partners and expressed them, either alone or together with their strongly biased Cys-less partners. In the weakly biased partners, we inserted the single Cys residue in a loop location that should be accessible to the extracellular (periplasmic) side if the protein is correctly oriented in an orientation that is opposite to that of the strongly biased partner (see location of the Cys residues in Fig. 1*B*, red circles). We then examined the orientations of the weakly biased monomer in the membrane by determining the location of the test cysteine using the SCAM method (<u>s</u>ubstituted <u>c</u>ys <u>a</u>ccessibility <u>m</u>ethod) (27, 28). In this assay, a sulfhydryl reagent that cannot cross the cytoplasmic membrane (we used AMS unless otherwise noted, see Materials and Methods) is added to whole cells to block any cysteines that are accessible from the periplasm. Subsequently, the membrane is disrupted and another reagent, maleimide-polyethylene glycol (mal-PEG), is added in order to derivatize any unblocked cysteines. Cytosolic cysteines remain reactive to mal-PEG, which increases the mass of the protein by ∼2 or ∼5 kDa, depending on the PEG size. In contrast, periplasmic cysteines are blocked by AMS and cannot subsequently be PEGylated. As a control, a membrane-permeable reagent, NEM, was used in the blocking step instead of AMS, to confirm that all test cysteines are in a solvent-exposed position that can be blocked. All proteins were specifically radiolabeled *in vivo* using the T7-rifamipicin method (27) in order to avoid the use of tags that may bias the topology.

For example, in the left four lanes in the top panel of Fig. 1*C*, lanes 1 and 4 show an unblocked protein with or without the PEGylation step, revealing the migration pattern of the un-PEGylated and PEGylated protein, respectively. In lane 2, pre-blocking by the membrane-permeable NEM was carried out prior to PEGylation and the blocking of PEGylation confirms that the introduced Cys is in an accessible position (either facing the cytosol or the periplasm). Finally, lane 3 shows that the membrane-impermeable reagent AMS blocks ∼70% of the PEGylation, indicating that the test Cys faces the periplasm in about 70% of the protein population.

All four subunits tested show high degrees of PEGylation blocking by AMS (70-100%, Fig. 1*C,D*), indicating periplasmic localization of the respective cysteines both when the monomers are expressed alone and together with their partner. The observed minor differences are probably due to selective degradation of un-complexed subunits (see below). This suggests that the weakly biased proteins are oriented properly already in the monomeric state, and that dimerization plays no role in achieving the final topology. Thus, their polypeptide sequence alone contains sufficient cues to orient them properly, even though the K/R-bias is weak. We could confirm that two of the protein pairs, Pairs #1 and #3, confer resistance against methyl viologen (Fig. S2). Notably, in both cases, both proteins of the pair (termed A and B) had to be coexpressed in order to confer resistance, confirming that a heterodimer is the functional unit. The introduced Cys mutations did not impair the function of the dimer in these two cases (Fig. S2).

Our finding that the weakly biased monomers are correctly oriented before dimerization suggests that these monomers do not undergo post-translational topological flip-flop in the membrane. As a potentially more sensitive test of this conclusion, we mutated the K/R-bias of an already weakly biased monomer, protein B from Pair #3, to bring the subunit towards a dual topology. Two cytoplasmic positive charges were mutated to alanines and an arginine was added at the periplasmic C-terminus, to generate the mutant K27A/R87A/R110 (Fig. 1*E*). The mutant was still functional in the methyl viologen resistance assay when coexpressed with its partner (Fig S2, lower panel), and had dual topology, showing only 45% periplasmic blocking of Cys109 located at the C-terminus when expressed alone (Fig. 1*F, G*). Coexpression of its partner (protein A) did not appreciably change the degree of blocking, again indicating that the B monomer does not undergo post-translational flip-flop, but that only the fraction already inserted in the orientation required to form anti-parallel dimers with the A monomer is available for dimerization.

Collectively, these results indicate that all subunits in heterodimeric SMR proteins orient correctly on their own – either by strong K/R-biases or by other less well-understood signals – and that dimerization with the binding partner is unlikely to play a significant role in determining the final orientation.

### Coexpression of topology-dedicated EmrE mutants does not affect the dual topology of EmrE

We next turned our attention to dual-topology homodimeric SMR members, in which identical monomers associate in opposite orientations to form the anti-parallel dimer. The best-studied example, EmrE from *E. coli*, was shown to exist as an anti-parallel dimer in the membrane with roughly 50% of the protomers having their termini outside of the cell (N_out_/C_out_) and 50 % inside (N_in_/C_in_) (22). We reasoned that post-translational topological flip-flop might be more common in dual-topology monomers, and may be important for the efficient production of anti-parallel dimers.

To follow the topology of EmrE, we engineered functional (Fig. S3*A*) single Cys mutations, either in the first loop (T28C) or at the C-terminus (position 111), pointing to different sides of the membrane (Fig. 2*A*). Both positions showed 50% periplasmic blocking of PEGylation by AMS, consistent with dual topology (Fig. 2*B,C*). As a control, we also carried out periplasmic blocking experiments for previously described mutants engineered to a single topology – either N_in_/C_in_ or N_out_/C_out_ (15, 16). The results are consistent with a single topology of these mutants (Fig. S3*B*).

**Figure 2.**
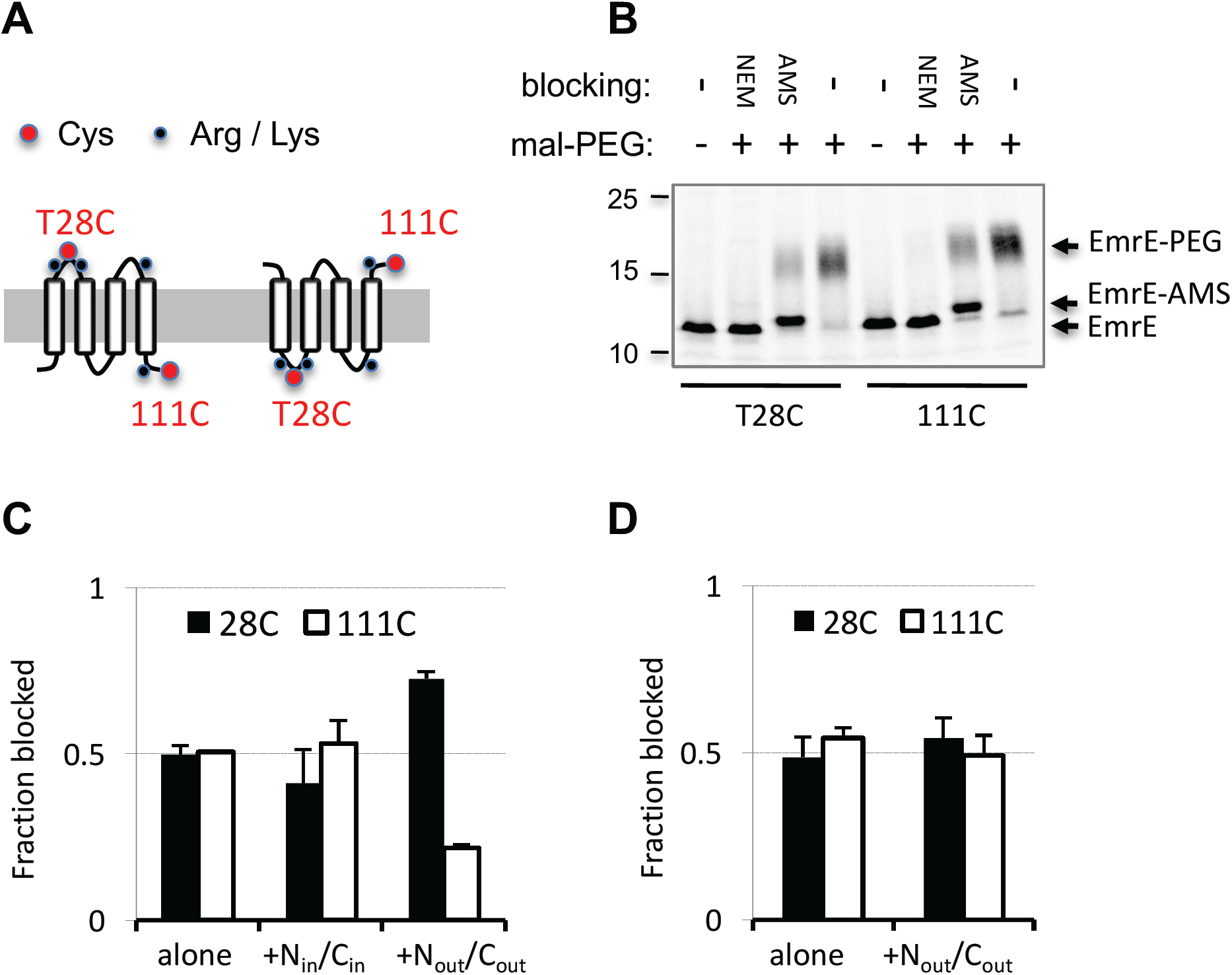
Coexpression of dual topology EmrE with topology-dedicated mutants. (A) Scheme showing the dual topology of EmrE, similar to Fig. 1*B*. The location of introduced single cysteine mutations is shown in red. Note that each mutant contains only a single Cys (either at position 28 or 111). (B) Example of SCAM of single Cys EmrE expressed alone, similar to Fig. 1*C*. The Cys mutant used is given below. (C) Quantified SCAM results for dual-topology EmrE (either T28C or 111C) expressed alone or with the N_in_/C_in_ or N_out_/C_out_ mutants. Error bars, SD of at least 3 independent experiments. (D) Quantified SCAM results for dual topology EmrE (either T28C or 111C) expressed alone or with the N_out_/C_out_ mutant in the Δ*ftsH* strain.

We utilized the same strategy as above to test if EmrE can flip-flop in the membrane, namely, we asked if expression of the engineered N_in_/C_in_ or N_out_/C_out_ mutants affected the final topology of a coexpressed dual-topology monomer. The results show that expression of EmrE alone or together with EmrE(N_in_/C_in_) results in roughly 50% blocking of the cysteines in positions 28 and 111, consistent with a 50:50 orientational distribution in both cases (Fig. 2*C*). In contrast, coexpression with EmrE(N_out_/C_out_) results in redistribution, biasing the wild-type monomer towards 75% N_in_/C_in_, with position 28 mainly in the periplasm (AMS-blocked) and position 111 mainly in the cytosol (protected). One explanation for this is that EmrE spontaneously reorients in the membrane to assemble with the surrounding unpaired N_out_/C_out_ subunits. An alternative explanation, however, is that EmrE(N_out_/C_out_) monomers compete with wild-type N_out_/C_out_ monomers for dimerization with the wild-type N_in_/C_in_ monomers, leaving a fraction of the wild-type N_out_/C_out_ monomers in an unpaired state possibly susceptible to rapid degradation. A higher fraction of N_in_/C_in_ than N_out_/C_out_ subunits would therefore remain in the membrane.

To test this scenario, we repeated the experiment in an *E. coli* strain deleted for the FtsH protease, which was recently shown to degrade un-complexed EmrE (13). Indeed, in this strain coexpression of the N_out_/C_out_ subunit no longer affected the topology of wild-type EmrE, with both T28C and C111 being blocked from the periplasm by AMS to about 50%, just as for wild-type EmrE expressed alone (Fig. 2*D*). Thus, degradation is likely the mechanism accounting for the observed biased topology upon coexpression. Taken together, these results indicate that wild-type EmrE monomers remain stably inserted in the membrane after the cotranslational insertion step.

### EmrE topology does not depend on dimerization

We devised a final strategy to examine if dimerization could affect the topology of EmrE monomers. We reasoned that if monomers could dynamically flip-flop in the membrane until they dimerize, this would have a buffering effect; namely that even if EmrE monomers were initially inserted in an un-balanced orientational distribution, anti-parallel dimerization would eventually result in a perfectly balanced 50-50 distribution. Accordingly, post-translational flip-flop, if possible, should make monomer orientation less sensitive to positive charge mutations that bias the initial insertion towards one topology or the other. Conversely, an EmrE mutant that cannot dimerize would flip back and forth randomly and would not benefit from this buffering effect. Therefore, the orientation of a non-dimerizing mutant should be more sensitive than wild-type EmrE to changes in the K/R-bias.

To study this, a non-dimerizing mutant was needed. The G97P mutation was previously shown to abolish EmrE dimerization (29). We confirmed this by two additional methods: (i) blue-native polyacrylamide electrophoresis (16), which showed that EmrE(G97P) runs primarily as a monomer, unlike the dimeric wild-type (Fig. 3*A*) and (ii) *in vivo* crosslinking of EmrE(S107C) (30), which confirmed that introducing the G97P mutation largely abolished the ability to crosslink the dimer of EmrE(S107C) in the native membrane (Fig. 3*B*).

**Figure 3.**
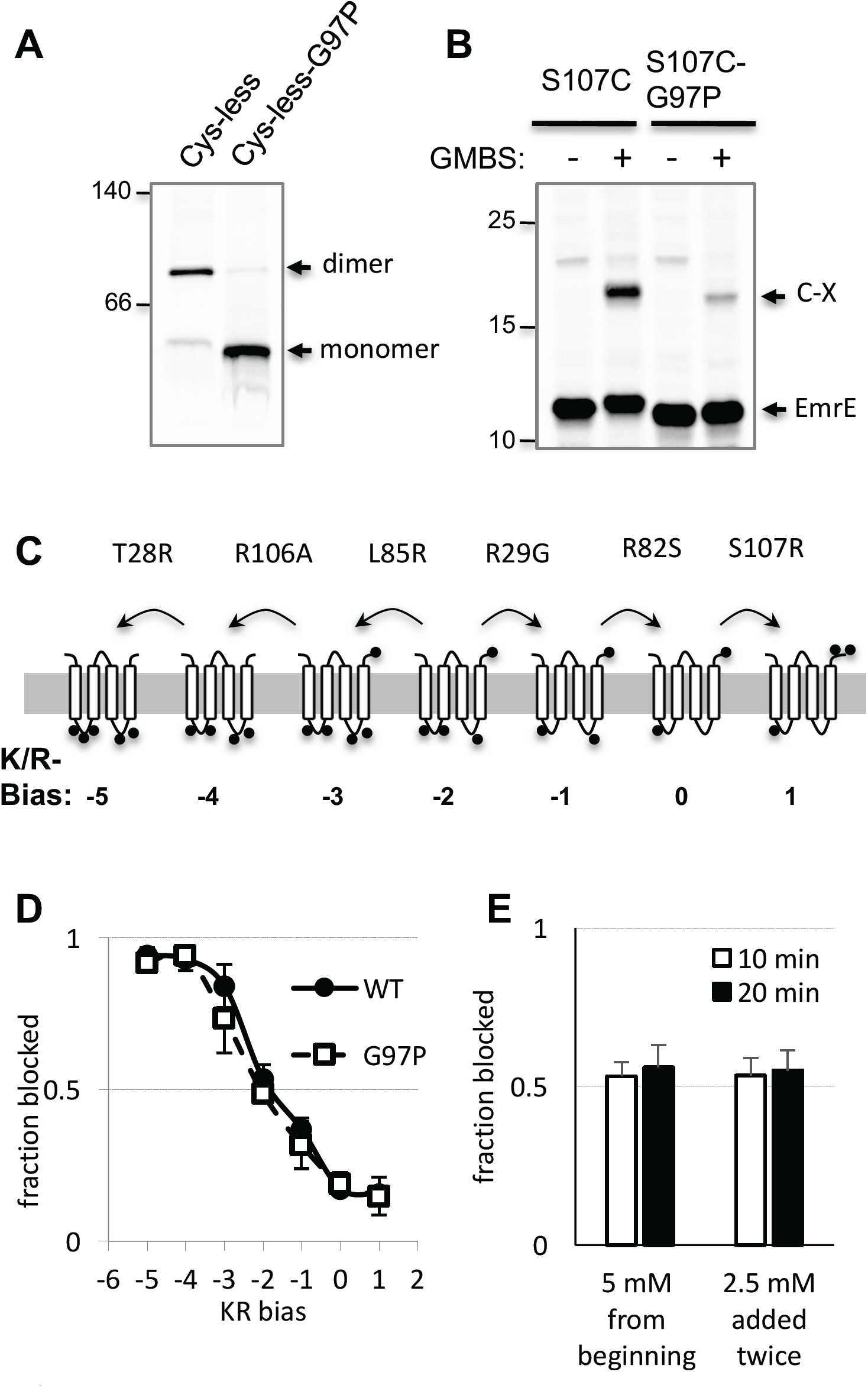
Dimerization does not affect the topology of EmrE mutants. (A) BN-PAGE of complexes formed by wild-type and mutant (G97P) EmrE. Migration of molecular weight native standard is shown to the left. *(B) In vivo* crosslinking of EmrE(S107C) and the G97P mutant by the reagent GMBS (N-γ-maleimidobutyryl-oxysuccinimide ester). C-X: crosslinked product. (C) Sequential positive charge mutants of EmrE on the path to N_in_/C_in_ (R29G-R82S-S107R, right) and N_out_/C_out_ topologies (L85R-R106A-T28C, left). Lysines and Arginines are indicated by black circles. The wild-type EmrE is represented by the middle protein. Note that in contrast to the depiction, the mutations affect EmrE orientation (D) Effect of charge mutations shown in (*C*) on the topology of EmrE and the G97P mutant, as inferred from SCAM of EmrE(111C). Error bars, SD (N = 3). (E) Time dependent blocking of EmrE(G97P-111C) following a single addition of 5 mM AMS from the beginning or two doses of 2.5 AMS mM separated by 10 min. Error bars, SD (N = 3).

Next, we tested the topology of a series of functional EmrE mutants, in which one, two, or three positive charges have been added or removed to bias the topology (see mutations in Fig. 3*C* (15)). Topology was determined by assessing the periplasmic accessibility of the Cys-111 mutant that was used above. The results show that the mutations act to gradually change the topology of EmrE, bringing Cys-111 to almost complete periplasmic or cytoplasmic exposure at the extreme K/R-biases (Fig. 3*D*). Starting from wild-type EmrE, two mutations were sufficient to bring the protein to a primarily single topology in either direction. Strikingly, the topology of the non-dimerizing EmrE(G97P) reacts identically to the charge mutations, within experimental error. Thus, the ability to dimerize does not impose a topological buffering effect on EmrE, suggesting that by the time EmrE subunits are available for dimerization, their topologies cannot change. Furthermore, the dual topology assumed by EmrE(G97P-111C) carrying no additional charge mutations (K/R-bias of −2 in Fig. 3*C,D*), even though it cannot dimerize, illustrates that dimerization is not required to reach a precisely matched 50-50 dual topology.

The 50% blocking of the monomeric EmrE(G97P-111C) further suggests that the monomer cannot flip-flop between N_in_/C_in_ and N_out_/C_out_ topologies on the time scale of our experiment (∼20 min), since if it could undergo rapid flip-flop, every monomer would spend half the time in the N_out_/C_out_ orientation. This would expose Cys-111 for periplasmic blocking by AMS in the entire protein population during the 20 minutes blocking reaction at 30°C. To ensure that the AMS concentration was not too low, nor that its reactivity diminished quickly over time, we carried out a control experiment in which half of the AMS was added initially and a second half was added after 10 min, increasing the time available for flip-flop. We then determined the level of periplasmic blocking after 10 or 20 minutes (Fig. 3*E*). The results show that 50% blocking was achieved already after 10 min, regardless of the AMS dosage, and that fresh AMS addition after 10 min did not lead to further blocking. Thus, the topology of individual monomers is stable within at most a few minutes after synthesis.

## Discussion

Understanding the degree of topological dynamics in fully synthesized membrane proteins is crucial for understanding their biogenesis. Yet, this problem is not easy to tackle. The membrane is a complex anisotropic environment. Its level of hydrophobicity changes gradually from the very hydrophilic aqueous solvent, through the lipid head-group region, and finally reaches maximal hydrophobicity in the internal hydrocarbon layer (2). From a physical chemistry standpoint, topological flip-flop may seem unlikely, or should at least be very slow. The loops that connect TMHs are decorated with charged groups that are not expected to be easily translocated across the membrane. Similarly, hydrophobic TMHs are expected to be quite stable in their trans-membrane disposition. Notably, though, not all loops and TMHs are created equal. Some TMHs are less hydrophobic than others and some loops are less charged, possibly reducing the energy barriers for topological changes. The *in vivo* situation is even more complex. Biological membranes are crowded with diverse proteins, some of which could potentially act as chaperones that may catalyze topological changes (2, 31). Only a few such chaperones/insertases have been described, and their mode of action is poorly understood.

With so many potentially contributing factors, it is not easy to predict how membrane proteins will behave *in vivo*. Several studies have suggested a surprisingly dynamic topology of membrane proteins under certain circumstances. In some proteins, it was observed that C-terminal parts of the protein can cause insertion or flipping of already inserted N-terminal parts of the protein (4, 5, 13, 11). More recently, it was suggested that EmrE can undergo wholesale flipping in the membrane following cleavage of a C-terminally-located stretch of positively charged residues (12). The underlying mechanism, however, is still obscure: Are membrane chaperones involved? Is topological dynamics only possible as long as the protein interacts with the membrane-insertion machinery? Or is topological dynamics possible only when starting from high-energy, topologically frustrated states where one or more TMH is not properly inserted into the membrane?

In the present study, we find no evidence for wholesale post-translational flip-flop of SMR monomers in the membrane. Instead, each individual monomer appears to attain a stable topology either during or immediately after translocon-mediated membrane insertion. In the case of dual-topology proteins such as *E. coli* EmrE, it thus appears that evolution has fine-tuned the polypeptide sequence such that the monomers insert stochastically with equal probability in either the N_in_/C_in_ or the N_out_/C_out_ orientation, rather than relying on a mechanism where the 50-50 distribution is reached only as a result of monomer flip-flop and subsequent formation of thermodynamically stable anti-parallel dimers.

Previous work has convincingly shown that N-terminally tagged EmrE monomers with an added, C-terminally located positively charged tail get initially inserted into the inner membrane in a frustrated N_out_/C_in_ topology. These monomers can then relax to the normal 4-TMH N_out_/C_out_ topology after cleavage of the C-terminal tail (12). Together with our observation that untagged EmrE N_in_/C_in_ and N_out_/C_out_ monomers do not undergo post-insertion topological flip-flop, this suggests a topological maturation scheme where individual EmrE monomers may initially insert in a mixture of different topologies, whereupon monomers with frustrated topologies (i.e., topologies where < 4 TMHs cross the membrane) rapidly relax to stable N_in_/C_in_ or N_out_/C_out_ topologies, much along the lines suggested by a recent molecular dynamics study (14). Of note, however, this model does not explain the apparent post-insertion flipping of N-terminally untagged EmrE monomers with 4-TMH N_in_/C_in_ topology suggested in the same study (12); further work will be required to resolve this discrepancy between the two sets of results.

SMR transporters are small membrane proteins and the extramembrane loops that connect their TMHs tend not to be very large or hydrophilic (12). Flip-flop in such proteins should therefore be fast compared to the majority of membrane proteins. The absence of flip-flop that we observe here thus suggests that wholesale reorientation is an unlikely event for proteins in biological membranes *in vivo*. Therefore, even small dual-topology proteins need to be initially inserted with an overall correct orientation.

## METHODS

### Bioinformatic analysis of SMR transporters

The UniprotKB (as of May 2013) was searched for the term COG2076, yielding 1015 SMR family genes, along with their protein sequences and Refseq identifiers. Entries for all proteins were retrieved in GenPept format from NCBI Protein database, from which the gene locations in their respective genome were taken. A multiple sequence alignment of all sequences was generated using Clustal Omega (32) and the TMHs of every sequence were determined by TMHMM 2.0 (33). Sequences were manually corrected based on the alignment to remove long N-terminal extensions (by re-assigning the initiator methionine). Aberrant sequences (misaligned or missing some of the transmembrane segments) were removed together with their genomic partners if they had such (see below), leaving 990 proteins. Consensus positions of the TMH boundaries were calculated based on the TMH bounds of all proteins in the alignment and the transmembrane K/R-bias was calculated using the consensus TMH bounds. Proteins having more Ks and Rs at the membrane side of the termini received a positive value for K/R-bias and *vice versa*. To analyze heterodimeric members, we utilized the observation that they occur in pairs (operons) in bacterial genomes (24). Protein pairs were identified based on them being at a distance of less than 3000 bp and encoded on the same strand in their respective genomes. Identical pairs were removed. The final non-redundant set included 112 pairs (224 proteins).

### Bacterial strains

*E. coli* MC1061 and DH5α were used for plasmid propagation. For most protein expression experiments, *E. coli* BL21 (DE3) Δ*emrE::kana*^*R*^, Δ*mdtJI::Cm*^*R*^, deleted for the *emrE* gene and for the *mdtJI* operon was used to avoid potential interactions between expressed proteins and chromosomally-encoded SMR transporters. The strain was generated by P1 transduction of the resistance alleles into BL21(DE3) as described (34). The Δ*emrE::kana*^*R*^ allele was from the Keio collection (35). The Δ*mdtJI::Cm*^*R*^ allele was first generated in *E. coli* BW25113 as described (35, 36) using primers that amplify a linear fragment from the pKD3 plasmid. The fwd and rev primers are GCTGAATTAAGCGAAAATTAAAATAATTCTCTTGCAGGAGAAGGACAATGatgggaattagccatggtcc and GCTGCCCGACAGCGCGGGCAGCGTCTTCATCAGGCAAGTTTCACCATGATtgtaggctggagctgcttcg, respectively.

The Δ*ftsH* strain AR5090 (DE3) was obtained by derivatizing the Δ*ftsH* strain AR5090 (37) with the DE3 element using the λDE3 Lysogenization Kit (Novagen).

### Plasmids

All proteins used were Cys-less versions of the original proteins. EmrE was expressed from pET-Duet-1 vector as described (15), typically from the second MCS. When two EmrE variants were coexpressed, the single-topology mutants were expressed in excess from MCS1 which supports higher expression (16). Heterodimeric SMR transporters operons were cloned into pET19b (Novagen). The vector original ribosome binding site and His tag were omitted. The cloned operons included the endogenous ribosome binding site, namely the 20 bp upstream of the first gene start codon. Standard PCR procedures were followed for all mutagenesis steps. For expression of a single protein without it’s binding partner, the binding partner’s open reading frame was deleted, leaving only the 20 bp upstream of the remaining protein start codon. Gene pairs #1-3 were synthesized by GenScript (NJ, USA). Gene pair #4 was amplified from *P. Mirabilis* genome. All endogenous Cys residues were replaced by Ser. The proteins identities are given by their UniprotKB identifiers as follows. Pair 1, protein A and B are *Lawsonia intracellularis* Q1MPU9 and Q1MPU8, respectively. Pair 2, *Desulfovibrio desulfuricans* B8J441 and B8J442. Pair 3, *Edwardsiella ictaluri* C5B9U6 and C5B9U5. Pair 4, *Proteus mirabilis* B4EVU5 and B4EVU6. The reporter Cys for SCAM analysis were as follows: M96C in Pair 1 protein A, C74 in Pair 2 A, A109C in Pair 3 B, A110C in pair 4 B.

### Drug resistance assay

A single colony from a fresh transformation of BL21(DE3) Δ*emrE::kana*^*R*^, Δ*mdtJI::Cm*^*R*^, harboring the indicated pET plasmids was grown over night in LB medium supplemented with 100 µg/ml ampicillin. The culture was then back diluted 1:50 into the same medium and grown at 37°C to mid-logarithmic phase (OD_600_ ∼0.5). Five 10-fold serial dilutions were then prepared from the cultures, with the highest density being of OD 0.1. The serial dilution was spotted (3.5 µL) on LB-agar-ampicillin plates with drugs and various additives, or on identical plates containing the additives but without the drug as control. The plates were allowed to grow for one or two nights at 37°C. For EmrE, the plates contained 30 mM bis-tris-propane pH 7.0 and the drug ethidium bromide (50 µg/mL). For Pair #1, the plates contained 0.05 mM IPTG and the drug methyl-viologen (50 µg/mL). For Pair #3, the plates contained 0.1 mM IPTG and the drug methyl-viologen (40 µg/mL).

### Cell growth and specific radiolabeling of tag-less SMR proteins

A single colony from a fresh transformation of BL21(DE3) Δ*emrE::kana*^*R*^, Δ*mdtJI::Cm*^*R*^ harboring plasmids encoding the proteins of interest was grown overnight in minimal medium (M9 salts, 1 mg/ml CSM amino acids minus methionine, 0.4% glycerol, 100 μg/ml thiamine, 2 mM MgSO4, 0.1 mM CaCl2, 100 μg/ml ampicillin). The cultures were back diluted into the same medium to an OD_600_ of 0.1 and grown at 37°C to mid-log phase (OD_600_ ∼0.5). The cultures were then induced by 0.1 mM IPTG for 10 min, followed by 15 min incubation with 0.2 mg/ml rifampicin, shaking at 37°C. Proteins were labeled with 15 µCi [^35^S]Met for 5 min, then mixed with a high excess (2 mM) of non-radioactive methionine and put on ice for 5 min to stop the reaction.

When using the Δ*ftsH* strain AR5090(DE3), all overnight cultures were grown at 30°C in LB or LB-agar supplemented with 100 µg/ml ampicillin and 1% glucose. For experiments, cultures were washed and diluted into M9 medium as above but with glucose replacing glycerol.

### Topology determination by cysteine labeling

The protocol was adapted from Nasie et al.(27). Radiolabeled cells (typically from 4 ml culture) were washed twice by centrifugation at 4°C, 3,200×*g*, 5 min and resuspension in 1.5 mL of Na-Mg buffer (150 mM NaCl, 30 mM Tris–HCl pH 7.5, 5 mM MgSO_4_, 1 mM TCEP (Tris(2-carboxyethyl)phosphine)). Cells were pelleted again and resuspended in 1 mL Na-Mg buffer and divided to four microcentrifuge tubes, each containing 200 µL. Ten µL water were added to tubes 1 and 4, tubes 2 and 3 received 20 µL of 0.2 M NEM and 10 µL of 100 mM AMS, respectively. Samples were spun down, mixed gently and incubated in 30°C for 20 min with gentle shaking, then transferred back to ice. Ice-cold Na-Mg buffer (0.7 mL) was then mixed with the cells and they were immediately washed twice by centrifugation at 4°C, 10,000×*g*, 2 min and resuspension in 0.9 ml of Na-Mg buffer. The cells were then pelleted again and resuspended in 150 µL of lysozyme buffer (150 mM NaCl, 30 mM Tris–HCl pH 8, 10 mM EDTA, 1 mM TCEP, cOmplete protease inhibitor (Roche), 1 mg/ml lysozyme) and frozen in −20°C for at least 20 min. Cells were disrupted by thawing at 25°C for 5 min followed by shaking at 37°C for 10 min. Then 0.9 ml of DNase solution (15 mM MgSO_4_, 10 µg/mL DNase I, 1 mM TCEP, 0.5 mM PMSF (Phenylmethanesulfonyl fluoride)) was added and the samples were allowed to shake at 37°C for 10 min before transferring them to ice. Crude membranes were collected by centrifugation at 4°C, 20,000×*g*, 20 min and the pellet was resuspended in 50 µL of Na buffer (150 mM NaCl, 30 mM Tris–HCl pH 7.5, 1 mM TCEP). Membrane proteins were solubilized and PEGylated by adding 7 µL of 10% DDM (β-D-dodecyl maltoside, Anatrace) and 13 µL of 27mM mal-PEG (Methoxypolyethylene glycol maleimide, either 5kDa from Sigma or 2 kDa from Nanocs, NY, depending on the protein). The samples were incubated with continuous mixing at 30°C for 1 h and the reaction was quenched by mixing with 17.5 µL 5x Sample Buffer (120 mM Tris–HCl pH 6.8, 50% glycerol, 100 mM DTT, 2% SDS, and 0.1% bromophenol blue). Samples were resolved on 15% SDS-PAGE gels run with a modified running buffer containing only 0.05% SDS (38). Gels were dried, visualized by autoradiography and quantified using an in-house custom software.

### Dimerization analysis by Blue-Native PAGE

Radiolabeled cell cultures (1.4 mL) were washed twice in Na-Mg buffer, disrupted by Lysozyme and crude membranes were prepared as described above for topology determination, except that 20 mM EDTA was added to the disrupted cells after the DNase treatment. The crude (20,000×*g*) membrane pellet was resuspended in 85 µL of ACA50 buffer (50 mM amino-n-caproic acid, 50 mM Bis-Tris, 0.5 mM EDTA, pH 7.0). Membranes were solubilized by incubating with 5 µL of 10% DDM for 1 h on ice, centrifuged for 1 min at 4°C, 20,000×*g* and 70 µL of the supernatant was transferred to ultracentrifuge tubes. Insoluble material was pelleted by centrifuging for 30 min at 4°C, 100,000×*g* and 30 µL of the supernatant was mixed with 30 µL of ACA750 buffer (750 mM amino-n-caproic acid, 50 mM Bis-Tris, 0.5 mM EDTA, pH 7.0) supplemented with 0.05% DDM. The samples were mixed with 5 µL of G250 solution (5% Coomassie G250 in ACA750 buffer) and loaded on a 13% BN-PAGE mini-gel (with a 5% stacking gel layer). Gels were run at 4°C at 100 V for 1 h, then 500 V. Gels were fixed, dried, and visualized by autoradiography.

### Cross-linking of EmrE dimers *in vivo*

The protocol was adapted from (30). Radiolabeled cell cultures (1.4 mL) were washed once in ice-cold 0.5 mL phosphate buffered saline (PBS), resuspended in 0.7 mL PBS and divided into 2 tubes, 0.3 mL each. The cells were mixed with 3 µL of freshly dissolved 5 mM GMBS (N-γ-maleimidobutyryl-oxysuccinimide ester, Thermo, in DMSO) or DMSO as control. Samples were incubated at 37°C for 20 min with gentle agitation and cross-linking was then quenched by incubating for 5 more min with 50 µL 1M Tris-HCl pH 7.5 at 37°C. Cells were pelleted and resuspended in 0.4 ml ice-cold PBS and TCA-precipitated using an equal volume of 20% ice-cold TCA. Acetone-washed pellets were dissolved in protein sample buffer (1x) supplemented with 80 mM dithiothreitol and 100 mM Tris-HCl pH 6.8 and heated to 80°C for 20 min. Ten µL samples were resolved on 12% NuPAGE Bis-Tris gel run with MES buffer (Invitrogen). Gels were fixed, dried, and visualized by autoradiography.

## Acknowledgements

We thank Harry L. T. Mobley (University of Michigan Medical School) for providing genomic DNA of *P. mirabilis*, Guenter Kramer (ZMBH, Heidelberg) for advice in operon construct design, Jan-Willem DeGier (Stockholm University) for providing several reagents and members of our group for helpful discussions. This work was supported by grants from the Knut and Alice Wallenberg Foundation, the Swedish Research Council, and the Swedish Cancer Foundation to GvH. NF was supported by long term postdoctoral fellowships from EMBO/Marie Curie Actions (ALTF 211-2014) and from HFSP (LT000277/2015-L).

---

**Figure S1.**
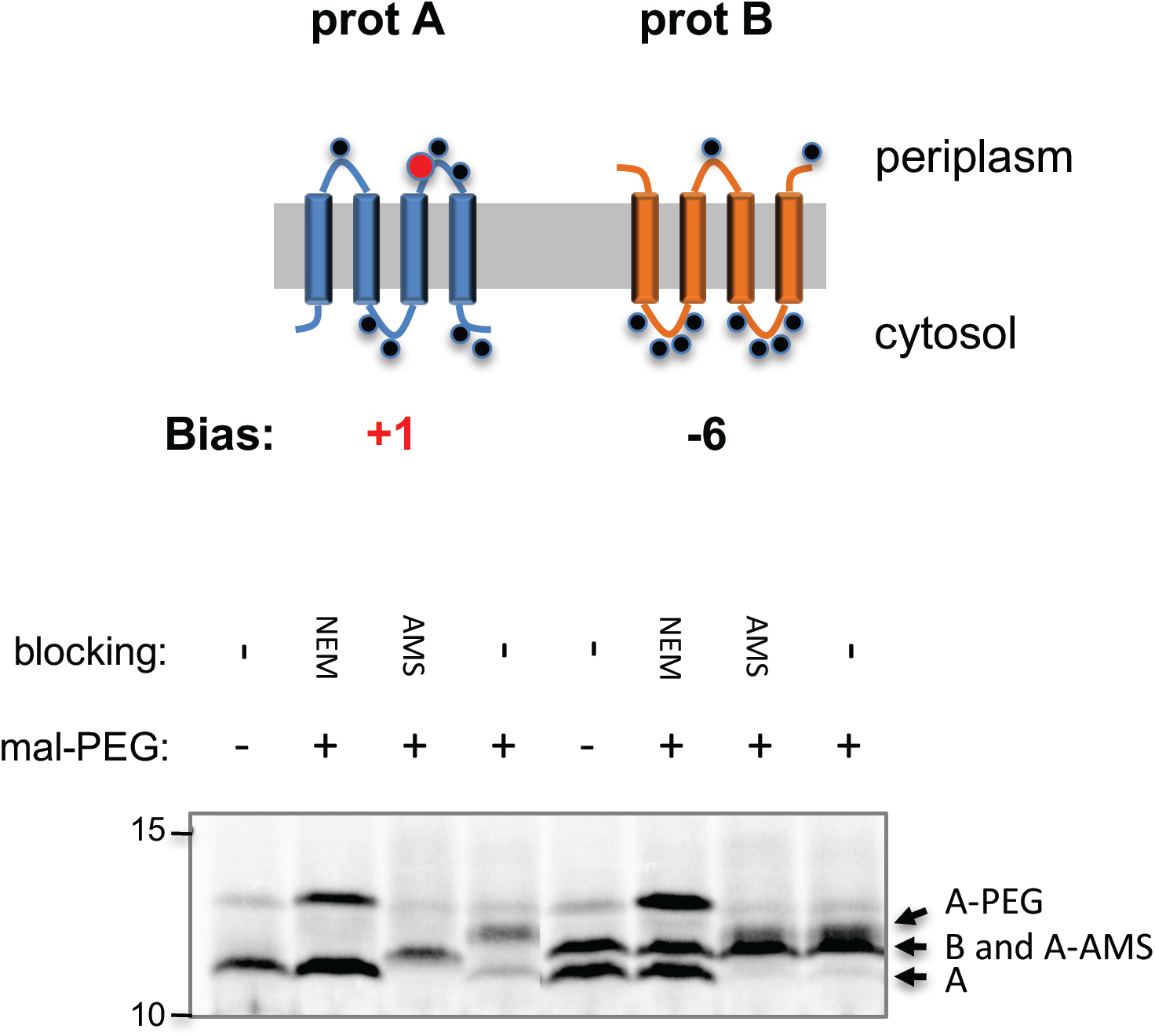
SCAM of heterodimeric pair #2 probed with AMS as the periplasmic blocker. See legend to Fig. 1*C*. The results indicate that residue Cys 74 of protein A (red circle in upper panel) is accessible to periplasmic AMS and blocked by it when protein A is expressed alone, but not together with it’s partner. This seems unlikely since the Cys is predicted to be accessible from the periplasm in the anti-parallel dimer (see model) and therefore expected to be blocked upon coexpression. Since topological changes seem an unlikely explanation for the difference in blocking, an alternative explanation is that local environment changes around Cys 74 occur due to interaction with protein B. Such changes may affect Cys 74 reactivity in the dimer. AMS is negatively charged; therefore one possibility is that in the heterodimer, Cys 74 could be in a negatively charged environment that repels AMS. A similar SCAM experiment using the positively charged reagent MTSET instead of AMS supports this. MTSET is also periplasmically-restricted, but can modify Cys 74 both in the monomer and the heterodimer (Fig. 1*C*)

**Figure S2.**
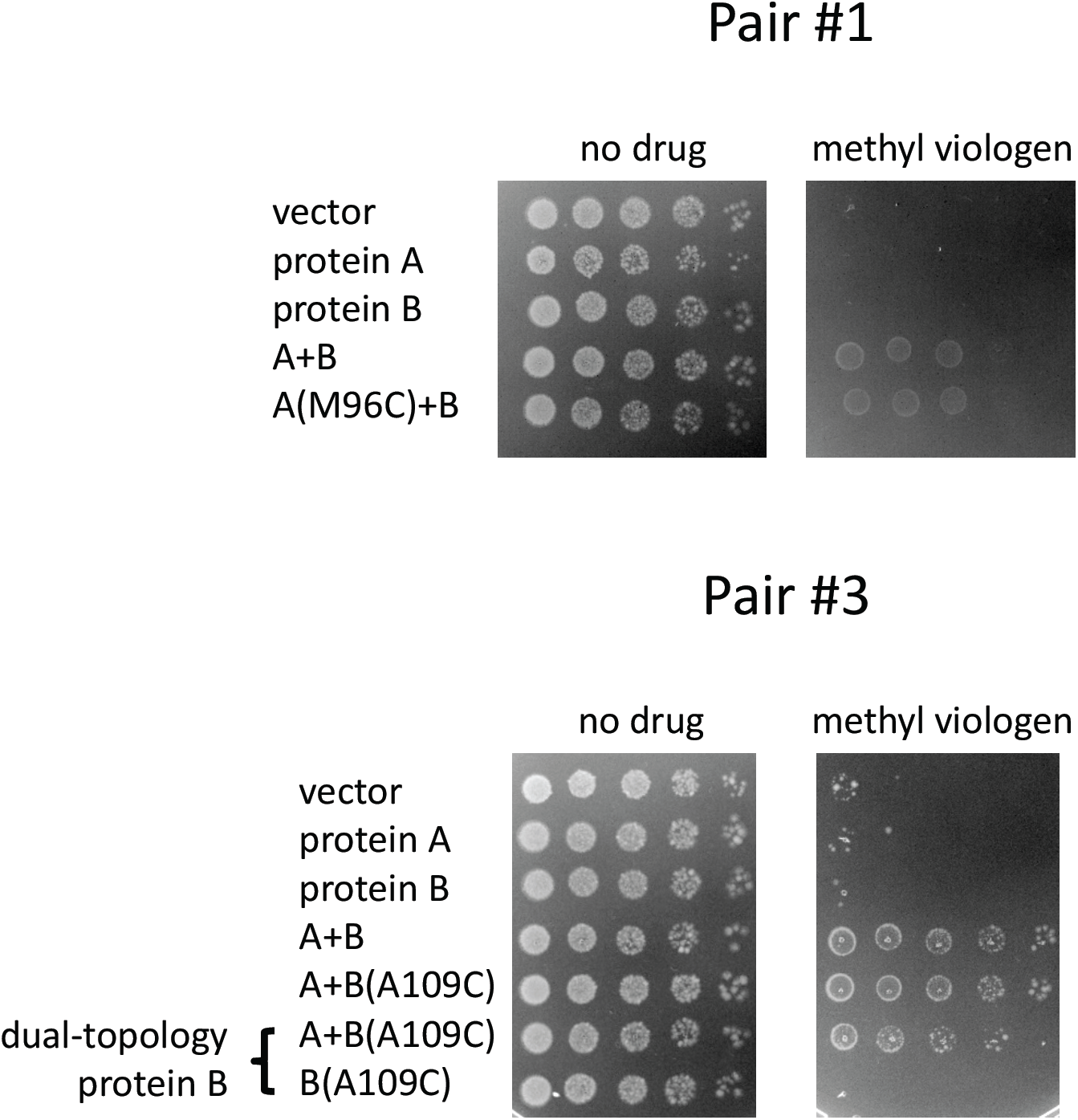
Methyl viologen resistance conferred by SMR transporter pairs mutants. A serial 10-fold dilution of *E. coli* BL21(DE3) Δ*emrE::kana*^*R*^, Δ*mdtJI::Cm*^*R*^ harboring plasmids containing the indicated constructs was spotted on plates with or without methyl-viologen. Cells harboring an empty vector serve as a negative control. The dual topology mutant of pair #3, protein B is K27A/R87A/R110. All proteins are Cys-less or single-Cys proteins, identical to those used for topology determination.

**Figure S3.**
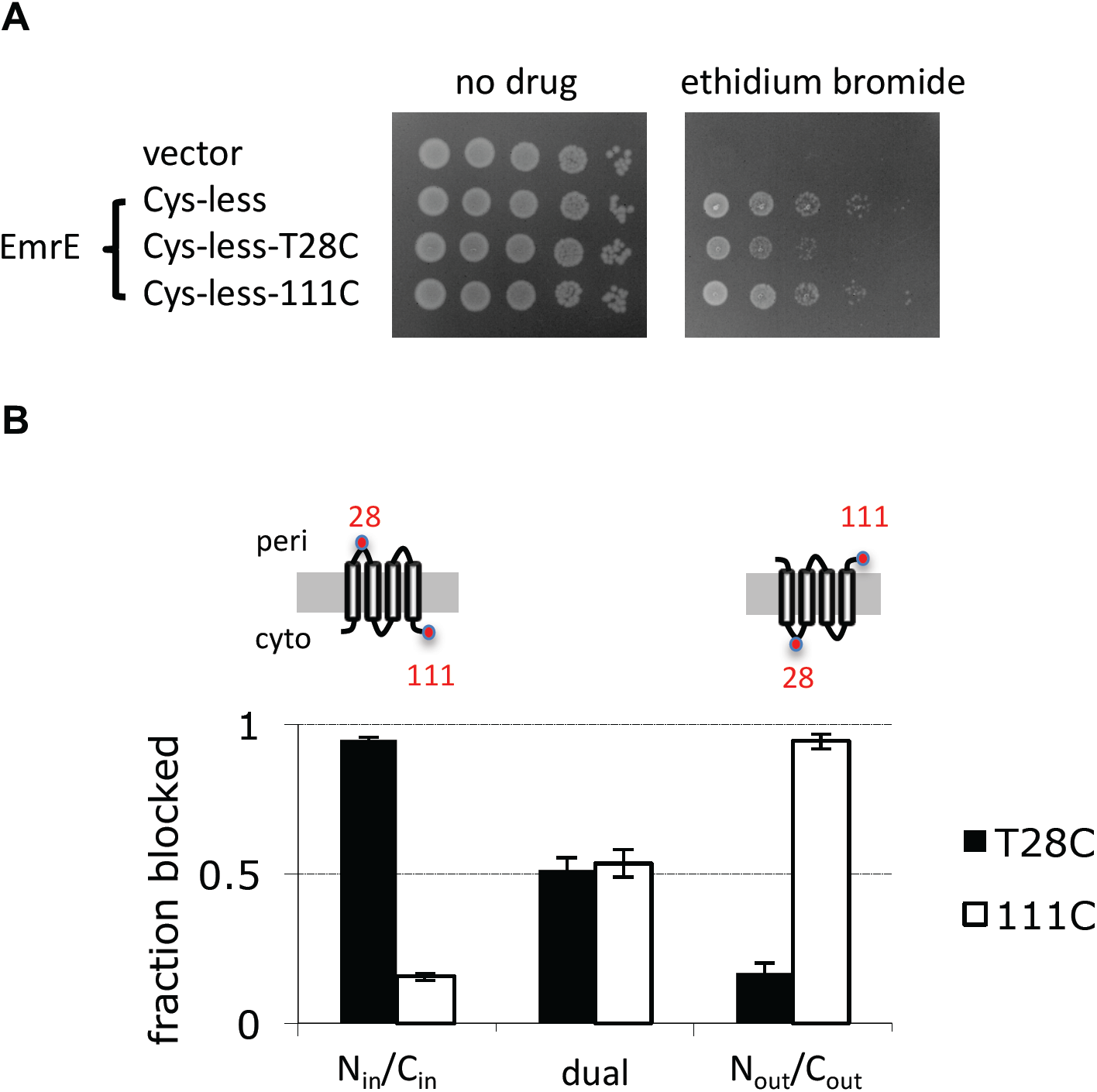
Characterization of Cys mutants engineered into different loops of EmrE. (A) The Cys mutants are functional, as inferred from their conferred ethidium bromide resistance. A serial 10-fold dilution of *E. coli* BL21(DE3) Δ*emrE::kana*^*R*^, Δ*mdtJI::Cm*^*R*^ harboring plasmids containing the indicated constructs was spotted on plates with or without ethidium bromide. Cells harboring an empty vector serve as a negative control. (B) Periplasmic blocking of the single Cys mutants in dual-topology EmrE and in single topology mutants engineered to N_in_/C_in_ or N_out_/C_out_. Note that N_out_/C_out_ EmrE(T28C) does not contain the mutation T28R from the original N_out_/C_out_ mutant, but two other charge mutations are included. The single topology mutants models are shown above their respective bars: the Cys pointing up (periplamic side) is expected to be blocked, while the Cys pointing down should not. Error bars, SD.

